# Feature Misbinding Underlying Serial-Order Effects of Visuospatial Working Memory

**DOI:** 10.1101/2025.07.03.657932

**Authors:** Linjing Jiang, Jennifer Tepan, Anastasiia Khibovska, Hoi-Chung Leung

## Abstract

The accurate processing of incoming visual information in serial order is fundamental to visual cognition. Prior studies have demonstrated primacy and recency effects in tasks requiring the serial recall of visual stimuli such as letters, digits, words, or locations. However, there is still ongoing debate about whether these primacy/recency effects on working memory retrieval reflect variable representational precision for multiple items in memory or misbinding of item features (e.g., location) and their ordinal position. This study sought to examine potential sources contributing to serial-position effects in visuospatial working memory using eye tracking and statistical modeling approaches. In two eye-tracking experiments, we measured the latency and endpoint error of serial-order memory-guided saccades under varying cue conditions (order cue vs. quadrant cue) from a total of 92 participants. The first memory-guided saccade (MGS) showed primacy and recency effects on endpoint error, latency, and transposition error in the order cue condition but not in the quadrant cue condition. Probabilistic modeling of MGS distribution showed a better fit of a standard (non-swapping) model to the quadrant-cue condition and a swap model to the order-cue condition. These findings indicate that visuospatial working memory representation varies across serial positions primarily due to location-serial-position misbinding rather than variable memory precision about location.

## Introduction

The ability to track the order of incoming sensory events is critical for many daily functions, ranging from verbal to visuospatial tasks. While serial-order verbal (visual and auditory) recall has been more extensively studied (See review by Hurlstone et al., 2014), relatively few researched the visuospatial domain (Bonanni et al., 2007; Farrand et al., 2001; Guérard & Tremblay, 2008; McAteer et al., 2023). It is unclear how exactly multiple items in a visuospatial sequence are represented in working memory and what latent cognitive sources contribute to variable serial-position memory effects. Using eye-tracking and statistical modeling, this study sought to address these issues, examining the nature of serial-order processing in visuospatial working memory.

Items in different serial positions are not remembered equally. A plethora of verbal short-term memory studies have demonstrated primacy and recency effects on response accuracy (e.g., Guérard & Saint-Aubin, 2012) and latency (e.g., Haberlandt et al., 2005) during serial recall of visual/auditory letters, digits, or words. Similar serial-position effects have been observed in visuospatial tasks, including serial reconstruction (Avons, 2007; Cortis et al., 2015; Hurlstone & Hitch, 2015; Jones et al., 1995; Martin et al., 2017), serial recall (Bonanni et al., 2007; Farrand et al., 2001; Guérard & Tremblay, 2008), and partial serial recall of spatial locations (Gorgoraptis et al., 2011; McAteer et al., 2023; Udale et al., 2022; Zokaei et al., 2011). For instance, the accuracy of recalling spatial locations in a sequence tends to decrease from early to intermediate items and then increase for last serial positions (e.g., Guérard & Tremblay, 2008). Interresponse latency, defined as the time between successive responses, declines sharply between the first and second serial positions and remains relatively stable thereafter (e.g., Hurlstone & Hitch, 2015).

Despite these consistent observations, the neurocognitive mechanisms underlying serial-position effects are not well understood. Features such as spatial location may be represented with varying precision across serial positions due to dynamic distribution of working memory resources among items (Gorgoraptis et al., 2011; Udale et al., 2022), as proposed in recent cognitive models of working memory such as the resource model (Bays et al., 2009; Bays & Husain, 2008). At the neural level, lower spatial working memory precision has been associated with higher variability of prefrontal neuronal activity during the delay period (Wimmer et al., 2014), suggesting a possible neural basis of variable spatial working memory precision across experimental factors such as serial position.

Additionally, serial-position effects may arise from the misbinding between visuospatial features and serial position of different objects in a sequence. Cognitive models of serial-order processing, such as the position-marking model (Brown et al., 2000) propose that items in a sequence are bound with contextual cues such as serial positions, while related working memory resource model (e.g., Bays et al., 2009; Manohar et al., 2019) suggest analogous misbinding for multi-feature stimuli in a group of memorized items. Such feature-serial-position binding may be achieved neurally by joint coding of spatial/visual and ordinal information (Ninokura et al., 2003, 2004) and via multiplicative gain modulation (Botvinick & Watanabe, 2007) in the lateral prefrontal cortex.

Statistical models, particularly mixture models (Bays et al., 2009), have been utilized to dissociate feature precision from misbinding to better interpret working memory data. However, findings from studies that model delayed recall in visual and spatial working memory across serial positions remain inconsistent. While some studies have reported significant serial-position effects on feature misbinding probability (Zokaei et al., 2011), others have found such effects on feature precision (McAteer et al., 2023) or on both (Gorgoraptis et al., 2011). Many of these studies introduced additional contextual cues, such as color, alongside spatial location and serial position (McAteer et al., 2023), complicating the interpretation of the misbinding effects. Moreover, the use of simultaneous presentation of items as a control condition, which eliminates the serial-position feature entirely, has limited the understanding of how visuospatial memory precision varies implicitly with serial positions.

In this study, we investigated serial-position effects and their cognitive source in visuospatial working memory using a novel eye-tracking task and statistical modeling. Across two experiments, a total of 92 participants completed a serial-order, memory-guided saccade task (sMGS) in which either the quadrant (quadrant cue condition) or the serial position (order cue condition) of a target was cued for recall. This manipulation made the order demand either implicit (quadrant cue) or explicit (order cue). To estimate location-serial-position misbinding probability and spatial memory precision, we applied various statistical models to analyze memory-guided saccade errors, latencies, and endpoint distributions across serial positions and cue conditions.

We hypothesized that serial-position effects are driven by both location-serial-position misbinding and changes in spatial working memory precision across memorized items. Specifically, we predicted that in the quadrant cue condition (implicit serial-position demand), recall error, latency, and memory precision would vary across serial positions, reflecting changes in spatial memory precision. The order cue condition (explicit serial-position demand) was expected to show stronger serial-position effects on recall error, latency, and swap rates compared to the quadrant cue condition, reflecting the additional demands of binding location to serial position.

Our design introduced several novelties: (1) The quadrant-cue condition allowed us to isolate the effects of spatial memory precision across serial positions without requiring explicit feature binding, as it required only the maintenance of serially presented locations. (2) The order-cue condition allowed us to specifically investigate the binding of location and serial position without additional visual features, such as color, often used in previous studies. (3) By measuring multiple types of eye movements and associated metrics during the response period, our approach afforded more refined behavioral measures than the manual-response methods commonly used in previous serial-position research.

## Experiment 1

Experiment 1 examined the effects of serial position on visuospatial working memory when order demand was manipulated within participants. Eye gaze positions were recorded from 65 participants while performing the sMGS task with the order cue and quadrant cue conditions as a within-subject factor. Error and latency of the primary and closest saccades to the target were analyzed and modeled across serial positions and cues.

### Participants

We recruited 65 college students from the Department of Psychology’s participant pool (age M = 20.4, SD = 3.6; 39 females). All participants had normal or corrected-to-normal vision and no self-reported neurological or psychiatric conditions. Informed consent was obtained from all participants. All the experimental procedures followed the protocol approved by the IRB committee at Stony Brook University. Course credits were given to the participants upon completion of the study session.

### Visual Stimuli

Visual stimuli used in the working memory task included eight images: two female faces, two male faces, and four houses. The face stimuli had neutral expressions and were selected from the NimStim database (Tottenham et al., 2009). The house stimuli were selected from the DalHouse database (Filliter et al., 2016). All stimuli were processed to yield the same resolution (500 x 500 pixels), mean, and standard deviation of luminance (CIE Lab, M = 48.7, SD = 24.3) using custom scripts in MATLAB and manual editing in Adobe Photoshop. The size of all image stimuli was 0.8 dva, and the size of the center cross and cues was 0.5 dva in diameter.

Visual stimuli were presented in 32 prototype locations (Fig. 1C). The eccentricity of the images was either 3 or 5.5 dva. The polar angle of the images was 15°, 35°, 55°, or 75° within each quadrant. Target locations were jittered across trials to reduce potential long-term memory effects, with the polar angle jittered between -5° to 5° and eccentricity jittered between -0.5 to 0.5 dva using a uniform distribution around the target.

**Figure 1.**
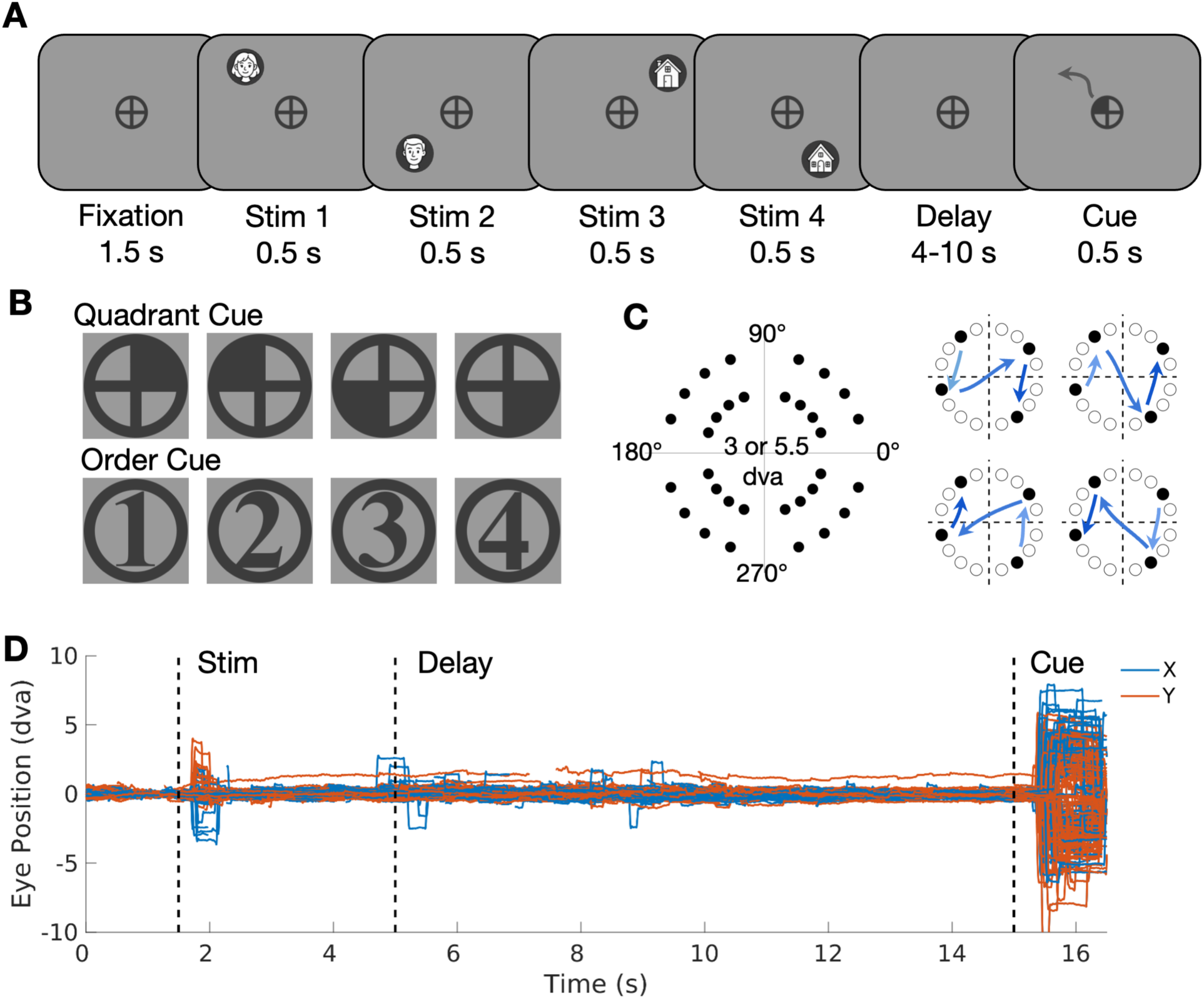
Serial-order Memory-Guided Saccade (sMGS) Task. (A) A schematic diagram of key events in a trial. Four items (a cartoon illustration of the stimuli is shown in the figure) were sequentially displayed in different locations on the screen with a 0.5-second interstimulus interval (not shown). After a delay of either 4, 6, 8, or 10 seconds, a task cue was shown to indicate the response target. Accordingly, the participants shift their eye gaze to the remembered target location. The gray curve illustrates a possible eye movement path during the response window (1 seconds), which was not presented in the actual task. (B) The two types of cues used in the experiment. During the quadrant cue condition, one quarter of the fixation stimulus turns black, indicating the response target’s quadrant. During the order cue condition, a number appears at the center of the fixation stimulus, indicating the response target’s serial position. (C) Target locations and sample sequence structures. There are 32 possible prototype locations, including 16 different polar angles (range between 15° to 345°) and two different eccentricities (3 and 5.5 degrees of visual angle, dva). On each trial, four items, one per quadrant at the same eccentricity, are presented sequentially either from left to right or from right to left to form four possible spatiotemporal patterns. Example spatial locations and the temporal order of the four items in a trial are displayed. (D) Sample gaze time course from all trials of a representative participant. Preprocessed eye gaze position in dva is plotted from fixation onset to the end of the response period. Horizontal (X) eye positions are plotted in red, and vertical (Y) eye positions are plotted in blue. Abbreviations: dva, degree of visual angle.

Tasks were programmed using Experimental Builder (SR Research, Ontario, Canada, Version 2.3.38) and displayed using a PC with Windows 10 operating system. The tasks were delivered on a Dell monitor with a resolution of 1280 x 1024 and a refresh rate of 75 Hz. Participants viewed the screen from a distance of 67.7 cm. The screen’s height was 30.5 cm, and its width was 38 cm, resulting in a maximum field of view of 31.4 degrees of visual angle (dva) horizontally and 25.4 dva vertically. Gaze positions were recorded monocularly at a sampling rate of 500 or 1000 Hz using a desktop-mounted, video-based eye tracker, the Eyelink 1000 (SR Research, Ontario, Canada).

### Task Design and Procedure

All participants completed a 1.5-hour study session consisting of 8 runs of the sMGS task. Each trial of the task (Fig. 1A) began with a 1.5-s fixation period, followed by the sequential presentation of four visual stimuli in different locations for 500 ms with a 500-ms inter-stimulus interval. Stimuli appeared at the same eccentricity (3 or 5.5 degrees of visual angle, dva), one in each quadrant. Following a delay of either 4, 6, 8, or 10 s, a cue (quadrant or order) appeared in the center of the screen, indicating which of the four remembered items is the response target (Fig. 1B). Specifically, the quadrant cue indicated the target quadrant by turning one-quarter of the fixation circle black, whereas the order cue indicated the serial position of the response target with a number displayed at the center of the fixation circle. Participants were instructed to shift their gaze to the remembered target location as quickly and accurately as possible within a 1.5-s response window. The inter-trial interval varied between 1.25, 1.5, and 1.75 s, average 1.5 s.

The main independent variables in this study were cue type and target serial position, which were manipulated within participants. Each participant completed four runs (64 trials) of the order cue task and four runs (64 trials) of the quadrant cue task, with the order of two tasks counterbalanced between participants. We probed all four serial positions for each cue condition an equal number of times.

In addition to the cue type and serial position, we controlled the spatiotemporal pattern of the sequence. The four items were presented in order from either the left to the right hemifield or from the right to the left hemifield in four temporal patterns (Fig. 1C). They appeared in 54 spatial forms (Fig. S1) at two levels of eccentricities (3 dva and 5.5 dva). The spatial forms of the sequences were carefully selected to be either fully asymmetric or have only one pair of dots symmetric relative to the x or y axis or the origin. These spatiotemporal patterns were designed to prevent subjects from predicting target location using sequence structure and to reduce both novelty and long-term memory effects.

The same spatiotemporal patterns were used for both cue conditions. Each of the four sequence patterns appeared four times in each run, and we probed the four serial positions for each of the four temporal patterns equally, resulting in 16 trials in total. Within a run, spatial forms were pseudo-randomly distributed across trials while meeting the following criteria: 1) Spatial patterns were maximally distinct across trials without repetition, based on a Procrustes analysis; 2) All 16 locations from one eccentricity appeared the same number of times; 3) All four quadrants were probed the same number of times. Each run included targets of a fixed eccentricity (e.g., 3 dva) and different eccentricities were counterbalanced between runs.

Lastly, we manipulated the order of the visual images (face first versus house first) within participants and the delay duration between participants (4/6 s: 30 participants; 8/10 s: 35 participants). The order of the images was counterbalanced between runs orthogonal to the eccentricities. After counterbalancing, we pseudo-randomly shuffled the trials to ensure the sequence was as unpredictable as possible.

### Oculomotor Analysis

Eye gaze data were preprocessed using customized MATLAB scripts. The raw data were smoothed using a second-order Savitzky–Golay filter with a kernel size of 20 ms (Nyström & Holmqvist, 2010). Artifacts such as blinks and noise were removed. Blinks were identified as samples with pupil velocity exceeding 8000 pixels per second, and an additional 100 ms of data before and after each blink were excluded to clean up surrounding noise. Samples with gaze positions outside the screen, gaze velocity greater than 1000 dva/s, or acceleration greater than 100,000 dva/s² (Nyström & Holmqvist, 2010) were identified as noise and removed without interpolation.

Following blink and artifact removal, an offline drift correction was performed (Vadillo et al., 2015). Baseline gaze positions were identified for each trial as fixations occurring 750 ms before stimulus onset. Fixations were defined as samples with a velocity ≤ 10 dva/s and duration ≥ 30 ms, and with both horizontal and vertical gaze dispersion ≤ 1.5 dva. For each trial, a linear transformation matrix was estimated as the mean distance between the mean baseline gaze location and the fixation cross location. This transformation matrix was applied to the horizontal and vertical gaze positions of all samples within that trial.

Saccades in the preprocessed data were classified as samples with a gaze velocity ≥ 30 dva/s, acceleration ≥ 6000 dva/s², amplitude ≥ 0.25 dva, and duration ≥ 8 ms. We defined several types of saccades for analysis. The first (primary) saccade within the 1500 ms response window after cue onset was analyzed to estimate participants’ initial memory recall on a trial-by-trial basis. Additionally, within the same time window, we identified the closest saccade to the target within 3.5 dva of the target and at least 2 dva from the fixation cross to estimate potential corrective saccades made to the target.

The endpoint error for both the first and closest saccades was calculated as the Euclidean distance between the mean position of the first fixation after the saccade and the target position in dva. The latency of the primary saccades was measured as the time between cue onset and saccade onset in ms.

Transposition errors were estimated by the percentage of trials in which participants responded to a non-cued stimulus. We calculated the polar angle of the primary saccade endpoint and compared it with the polar angle of the target. If the primary saccade landed within a different quadrant from the target, that trial was defined as having a transposition error.

All participants were included in the final analysis. We excluded the following types of trials from the final analysis: 1) There was a blink or an artifact during the first 500 ms of the response window (9.0% ± 9.9% of trials on average across participants); 2) There was no measurable baseline (0.8% ± 1.1% of trials); 3) There was no saccade detected during the response window (4.7% ± 6.4% of trials); 4) First saccade latency was less than 50 ms (0.08% ± 0.34% of trials); 5) First saccade start position was larger than 2 dva from the center cross (2.8% ± 3.5% of trials); 6) There was no fixation detected after the primary saccade (1.6% ± 3.4% of trials); 7) Other data collection error, such as missing pupil (1.0% ± 4.6% of trials).

After trial exclusion, on average 80.0% ± 14.0% of trials were included in the final analysis across participants. Among these trials, 65.7% ± 15.6% of trials were identified as responses to the target as their first saccade endpoint is less than 3.5 dva from the target location, 5.0% ± 3.2% of trials as responses to non-target (a saccade endpoint that is more than 3.5 dva from the target but less than 3.5 dva from one or more non-targets), 9.4% ± 6.7% of trials as responses outside of all regions of interest (a saccade endpoint more than 3.5 dva from any target/non-targets). The percentage of trials included for the quadrant cue condition (80.1% ± 14.5%) did not significantly differ from the order cue condition (80.0% ± 16.1%; *t*(64) = 0.06, *p* = 0.95).

### Model Design and Fitting

We used three two-dimensional probability mixture models (MemFit2D) (Grogan et al., 2020) to estimate sources of serial-order processing in spatial working memory. With the standard mixture model (Zhang & Luck, 2008), we estimated the probability of responses, 𝑝(𝑥), by a mixture of two distributions. Memory precision about the target was estimated by a bivariate Gaussian distribution centered around the target location 𝘛 with variance 𝜎^2^ (𝜎_𝑥_^2^ = 𝜎_𝑦_^2^ = 𝜎^2^, 𝜎_*xy*_^2^ = 0). A bivariate uniform distribution spanning the area of the entire screen, 𝐴, was estimated to account for random guessing. These distributions were weighted by 𝛼 and 𝛾 respectively, where 𝛼 + 𝛾 = 1 and 0 ≦ 𝛼, 𝛾 ≦ 1.

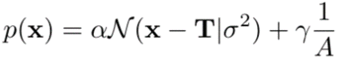

In the swap model (Bays et al., 2009), we also estimated response directed toward the non-targets (‘Swapping’) by including three bivariate Gaussian distributions (𝑚 = 3), centered around the three non-target locations 𝘕𝘛_𝑖_ with the same mean and variance as the target distribution. These distributions were weighted by 𝛼, 𝛽/𝑚, and 𝛾 respectively, where 𝛼+𝛽+𝛾=1 and 0 ≦ 𝛼, 𝛽, 𝛾 ≦ 1.

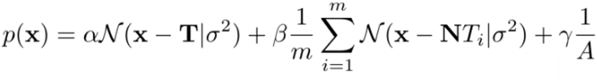

Lastly, in the swap plus systematic bias model (Grogan et al., 2020), we varied the mean of the bivariate target distribution (μ_*x*_ = μ_*y*_ = μ) to estimate the systematic bias of the responses in addition to memory precision.

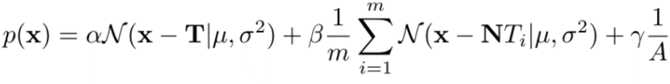

We fitted the models to the data using maximum likelihood estimation and derived the following parameters: 𝛼, probability of responding to target; 𝛽, probability of responding to non-target (swapping or feature misbinding); 𝛾, probability of random guessing; μ, systematic bias of responses; and 𝜎, memory imprecision (reciprocal of memory precision).

Model fitting was conducted on 13.3 trials on average (ranging from 3 to 16 trials) per serial position per cue type for each participant. The Akaike information criterion (AIC) (Akaike, 1974) and Bayesian information criterion (BIC) (Schwarz, 1978) were used to assess the goodness of fitting. Lower AIC or BIC values indicate better-fitting models. A difference in AIC or BIC values greater than 2 between models provides substantial evidence favoring the model with the lower value (Burnham & Anderson, 2004; Raftery, 1995).

## Statistical Analysis

Two-way repeated measures ANOVAs with serial position and cue were conducted on endpoint error and latency of the primary saccade and the closest saccades to the target. Additionally, modeled outputs, including target probability (𝛼), swapping rate (𝛽), guessing rate (𝛾), and memory imprecision (𝜎) were also analyzed across serial position and cue types.

## Results

### Memory-guided saccade error across serial positions

To examine how endpoint errors of memory-guided saccade vary by serial positions across cues, we conducted two-way repeated measures ANOVAs on the primary and closest saccades to the response target with serial position and cue type as within-subject factors (Fig. 2A, B).

**Figure 2.**
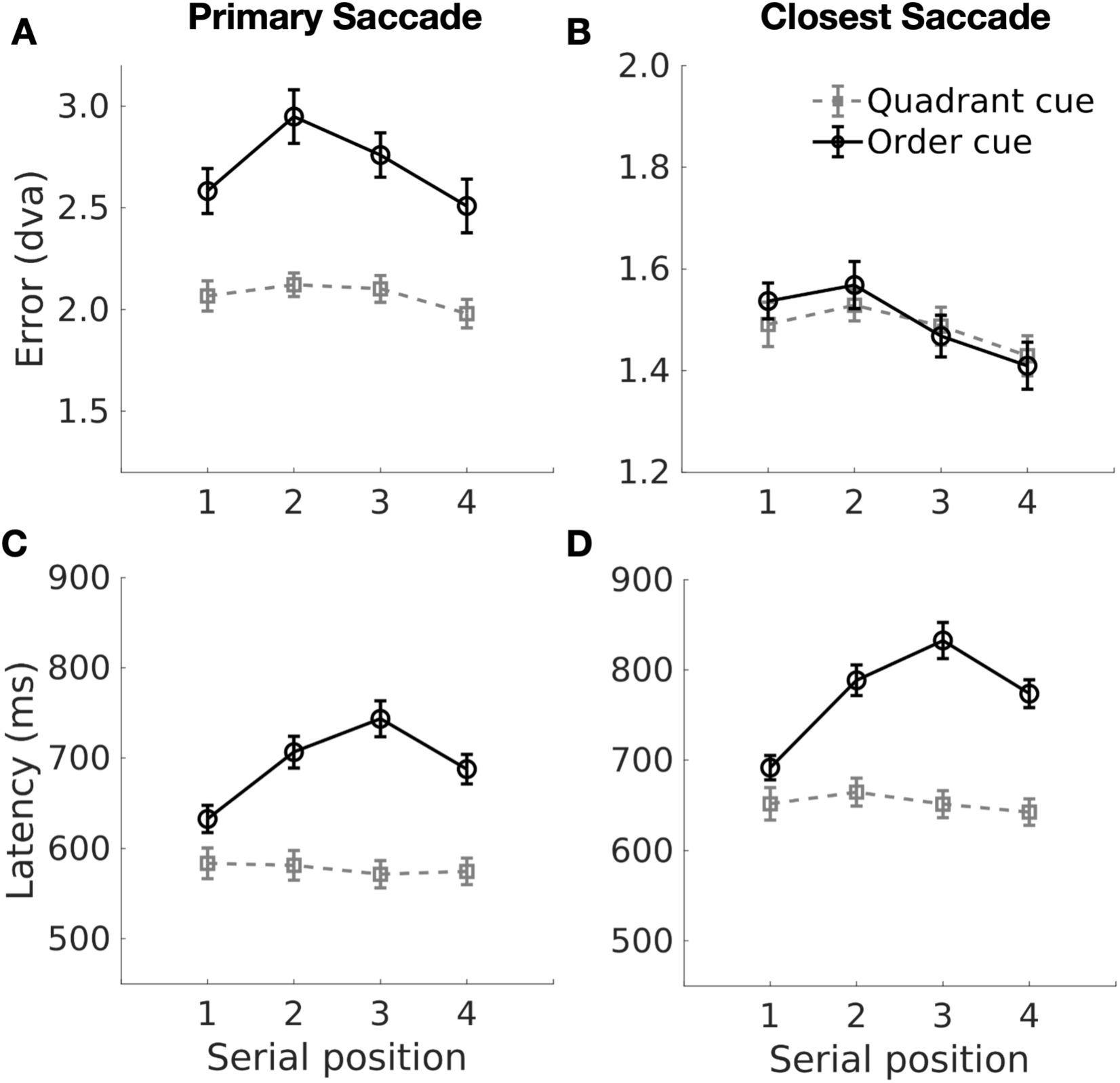
Memory-guided saccade endpoint errors across cue type and serial position in Experiment 1. Plots in the top row show saccade endpoint error for the two cue conditions at each serial position for the primary saccade (A) and the saccade that was closest to the target within the area of interest (B). Plots in the bottom row show the corresponding saccade latency (C, D). Y axes represent the saccade measures and X axes represent the serial position of the target in the stimulus sequence. Quadrant cue is plotted in gray and rectangle symbol; order cue is plotted in black and circle symbol.

For primary saccadic responses after cue presentation (Fig. 2A), we observed significant effects of cue (*F*(1, 64) = 60.40, *p* < 0.001, 𝜂_p_^2^ = 0.49), serial position (*F*(3, 192) = 9.36, *p* < 0.001, 𝜂_p_^2^ = 0.13), and their interaction (*F*(3, 192) = 3.05, *p* = 0.03, 𝜂_p_^2^ = 0.05). The interaction seemed to be driven by a significant main effect of serial position for the order cue condition (*F*(3, 192) = 7.56, *p* < 0.001, 𝜂_p_^2^ = 0.11) but not the quadrant cue condition (*F*(3, 192) = 2.25, *p* = 0.08, 𝜂_p_^2^ = 0.03). Polynomial contrast analysis of the order cue condition revealed a significant quadratic (*t*(192) = -4.32, *p* < 0.001) but not linear (*t*(192) = -1.28, *p* = 0.20) serial-position effect on saccade endpoint errors. Pairwise comparisons of serial positions revealed significant differences between positions one and two (*t*(64) = -3.63, *p* = 0.002, cohen’s *d* = -0.37) and between positions two and four (*t*(64) = 4.35, *p* < 0.001, cohen’s *d* = 0.45).

In contrast, endpoint error of the closest saccades toward the target (Fig. 2B) was significantly modulated by serial position (*F*(3, 192) = 4.81, *p* = 0.003, 𝜂_p_^2^ = 0.07) but not by cue type (*F*(1, 64) = 0.29, *p* = 0.59, 𝜂_p_^2^ = 0.004) and there was no significant interaction between them (*F*(3, 192) = 0.61, *p* = 0.61, 𝜂_p_^2^ = 0.009). Simple main effects of serial position were significant for the order cue (*F*(3, 192) = 3.68, *p* = 0.013, 𝜂_p_^2^ = 0.05) but not the quadrant cue condition (*F*(3, 192) = 1.72, *p* = 0.16, 𝜂_p_^2^ = 0.03). During the order cue condition, serial-position effects demonstrated a more linear (*t*(192) = -2.91, *p* = 0.004) and less quadratic (*t*(192) = -1.21, *p* = 0.23) trend with significant endpoint error differences between serial positions two and four (*t*(64) = 3.03, *p* = 0.017, Cohen’s *d* = 0.46).

Together, these results showed both primacy and recency effects on the first memory-guided saccades and a recency effect on the closest saccade to the target during the order cue condition. Memory-guided saccade errors during the quadrant cue condition, however, did not significantly vary by serial position.

### Memory-guided saccade latency across serial positions

To examine how latency of memory-guided saccade varies by serial positions across cue conditions, we conducted similar two-way repeated measures ANOVAs on saccade latency (Fig. 2C, D). For primary saccades (Fig. 2C), significant effects were found for serial position (*F*(2.6, 166.6) = 20.78, *p* < 0.001, 𝜂_p_^2^ = 0.25), cue type (*F*(1, 64) = 217.78, *p* < 0.001, 𝜂_p_^2^ = 0.77), and their interaction (*F*(3, 192) = 25.75, *p* < 0.001, 𝜂_p_^2^ = 0.29). The interaction was driven by the serial-position effect in the order cue condition (*F*(3, 192) = 38.91, *p* < 0.001, 𝜂_p_^2^ = 0.38), with saccade latency showing both linear (*t*(192) = 6.11, *p* < 0.001) and quadratic trends (*t*(192) = - 8.75, *p* < 0.001). During the order cue condition, all pairwise comparisons between serial positions reached statistical significance (*p*’s <= 0.001) except for the difference between serial positions two and four (*p* = 0.08). There was no significant serial-position effect in the quadrant cue condition (*F*(3, 192) = 0.86, *p* = 0.46, 𝜂_p_^2^ = 0.01).

Similar effects were observed for closest saccades to the response target (Fig. 2D), including serial position (*F*(2.7, 175.8) = 24.69, *p* < 0.001, 𝜂_p_^2^ = 0.28), cue type (*F*(1, 64) = 175.33, *p* < 0.001, 𝜂_p_^2^ = 0.73), and their interaction (*F*(3, 192) = 21.74, *p* < 0.001, 𝜂_p_^2^ = 0.25). Latency significantly (*F*(3, 192) = 36.77, *p* < 0.001, 𝜂_p_^2^ = 0.37) varied by serial positions during the order cue condition both linearly (*t*(192) = 6.68, *p* < 0.001) and quadratically (*t*(192) = -8.02, *p* < 0.001). During the order cue condition, all pairwise comparisons between serial positions were significant (*p* < 0.005) except between positions two and four (*p* = 0.28). There was no significant serial-position effect in the quadrant cue condition (*F*(3, 192) = 1.42, *p* = 0.24, 𝜂_p_^2^ = 0.02).

Together, the latency of both the primary and closest memory-guided saccades to target showed a primacy and recency effect during the order cue condition but no serial-position effects during the quadrant cue condition.

### Modeled memory precision, swap rate, and transposition errors across serial positions

To characterize the cognitive sources of serial-position effects on memory-guided saccades, we fitted three statistical models to the endpoint distribution of primary saccades.

The standard mixture model, incorporating only target and guessing distributions, best fitted the quadrant-cue data (Table 1). The order-cue data was best fitted by the swap model with the additional non-target distributions included in the model. Model fit was evaluated using AIC and by the standard mixture model according to BIC (Table 1). These results indicate increased item swapping in the order-cue condition.

**Table 1.**
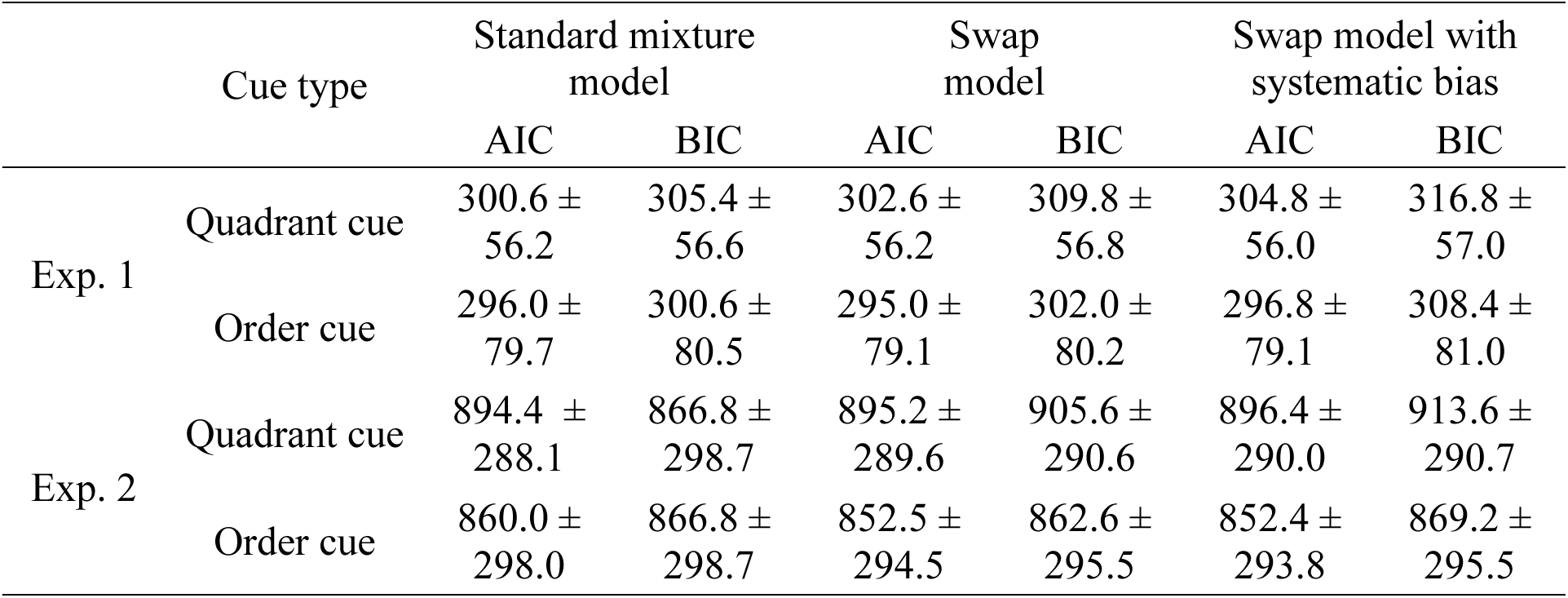
Model Fit for Experiments 1 and 2. AIC: Akaike Information Criterion; BIC: Bayesian Information Criterion.

The swap model was then used to estimate parameters including memory imprecision, swap rate, guessing rate, and target rate for the two cue conditions and serial positions. There was a significant effect of cue type on memory imprecision (Fig. 3A; *F*(1, 64) = 9.82, *p* = 0.003, 𝜂_p_^2^ = 0.13). Similar to the unmodeled data described above, the order-cue responses showed lower modeled memory precision (i.e., larger endpoint error) than the quadrant-cue responses (*t*(64) = 3.13, *p* = 0.003, Cohen’s *d* = 0.27). Effects of serial position on memory imprecision approached statistical significance (*F*(3, 192) = 2.32, *p* = 0.076, 𝜂_p_^2^ = 0.04). There was no interaction between cue type and serial position (*F*(3, 192) = 0.31, *p* = 0.82, 𝜂_p_^2^ = 0.005). Simple main effects of serial position did not reach statistical significance during both the quadrant cue (*F*(3, 192) = 2.05, *p* = 0.11, 𝜂_p_^2^ = 0.03) and the order cue condition (*F*(3, 192) = 0.92, *p* = 0.43, 𝜂_p_^2^ = 0.01).

**Figure 3.**
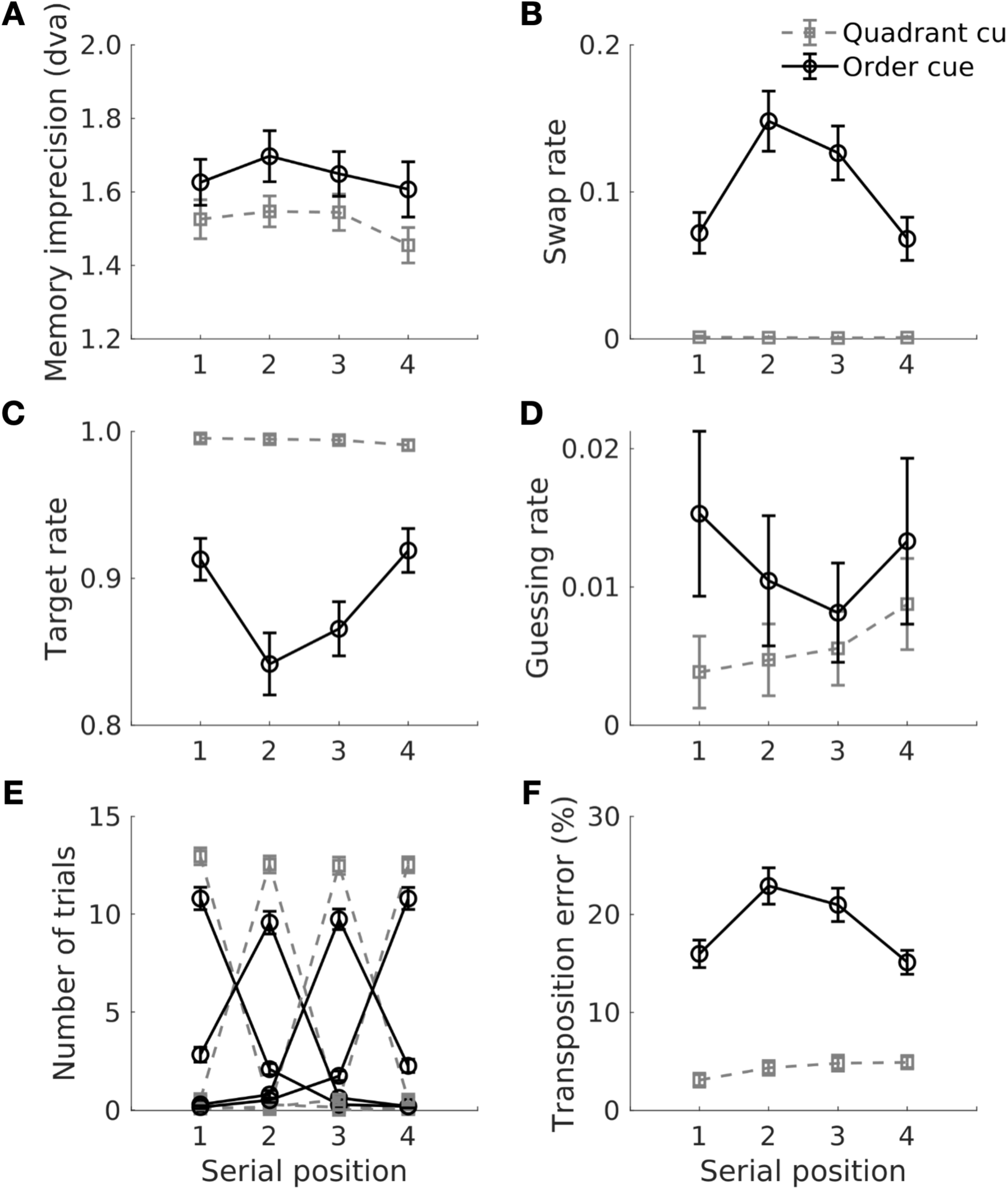
The Swap Model Results and Transposition Error of Memory-Guided Saccades in Experiment 1. (A) Modeled memory imprecision (standard error) in dva. Black and gray lines respectively represent the order and quadrant cue condition. (B) Modeled swap or misbinding rate (beta). (C) Modeled target probability (alpha). (D) Modeled guessing rate (gamma). (E) Number of trials (group average) during which the saccadic response was initiated toward the different serial positions. Lines of the same color represent trials of different target serial positions for the same cue condition. If the responses were always toward the correct item position, we would expect to see four clear peaks centered around each serial position as observed for the quadrant cue condition (gray lines). During the order cue condition, some of the saccadic responses were towards the non-targets (black lines) with lower peaks and higher valleys relative to the gray lines. (F) Transposition error, or the percentage of trials that included saccades responded to the non-target quadrant obtained from the empirical data.

For guessing rate (Fig. 3D), only the effect of cue reached statistical significance (*F*(1,64) = 4.95, *p* = 0.03, 𝜂_p_^2^ = 0.07), with a higher guessing rate for the order cue than the quadrant cue (*t*(64) = 2.23, *p* = 0.03, Cohen’s *d* = 0.18).

Swapping rate (Fig. 3B) significantly differed by the cue (*F*(1, 64) = 74.6, *p* < 0.001, 𝜂_p_^2^ = 0.54) and was higher in the order cue condition compared to the quadrant cue condition (*t*(64) = 8.64, *p* < 0.001, Cohen’s *d* = 1.06). There was a significant serial-position effect (*F*(3, 192) = 7.85, *p* < 0.001, 𝜂_p_^2^ = 0.11) and interaction between cue and serial position (*F*(3, 192) = 8.02, *p* < 0.001, 𝜂_p_^2^ = 0.11). That is, the swap rate varied significantly with serial position during the order cue (*F*(3, 192) = 7.95, *p* < 0.001, 𝜂_p_^2^ = 0.11) with a quadratic trend (*t*(192) = -4.76, *p* < 0.001), but the swap rate did not vary with serial position during the quadrant cue condition (*F*(3, 192) = 0.13, *p* = 0.94, 𝜂_p_^2^ = 0.002).

Consistent with the modeled swapping rate, transposition errors of the primary saccades (Fig. 3E, F) showed a significant effect of cue (*F*(1, 64) = 144.92, *p* < 0.001, 𝜂_p_^2^ = 0.69), serial position (*F*(3, 192) = 8.31, *p* < 0.001, 𝜂_p_^2^ = 0.12), and their interaction (*F*(3, 192) = 6.67, *p* < 0.001, 𝜂_p_^2^ = 0.09). Particularly, transposition errors varied significantly by serial position during the order cue condition (*F*(3, 192) = 9.69, *p* < 0.001, 𝜂_p_^2^ = 0.13) quadratically (*t*(192) = -5.25, p < 0.001) but not during the quadrant cue condition (*F*(3, 192) = 1.33, *p* = 0.27, 𝜂_p_^2^ = 0.02).

In sum, our results demonstrated a significant effect of cue type on memory-guided saccades. The order cue condition, compared to the quadrant cue condition, resulted in higher saccade endpoint error, saccade latency, transposition error, modeled swapping rate, and guessing rate, as well as lower modeled memory precision, showing an overall poorer performance with the explicit order demand. The serial-position effects were only evident under the order cue condition, with significant primacy and recency effects on the primary saccade endpoint error, latency, transposition error, and modeled swapping rate of the primary saccades. Additionally, a small recency effect was observed on the closest saccade to the target, and there was no significant serial-position effect on the modeled memory precision. These findings suggest that serial-position effects (both primacy and recency) occur only when order demand is explicit, and these effects may arise from the misbinding of spatial locations and serial positions rather than changes in memory precision.

## Experiment 2

One issue with Experiment 1 is its relatively small number of trials per serial position for each cue condition, which can affect model fitting. To further validate that memory misbinding as the major source of serial-position effects, we conducted a second experiment that collected more trials from a separate group of participants. In this experiment, we manipulated cue conditions between participants and increased the number of trials to up to 64 per serial position per cue condition for better modeling of the saccadic responses.

### Participants

27 participants were recruited from the Psychology Department’s subject pool (age M = 20.5, SD = 4.1, 16 females) for Experiment 2. All participants had normal or corrected-to-normal vision and no self-reported neurological or psychiatric conditions. Informed consent was obtained from all participants, and the experimental procedure followed the same protocol approved by the IRB committee at Stony Brook University as in Experiment 1.

### Task Design and Procedure

The task design and stimuli were similar to Experiment 1, with only the following differences. First, the delay duration was reduced to 4 or 6 s, as a supplementary analysis of Experiment 1 showed minimal effects of delay on task performance (Fig. S2). Second, the cue type (quadrant or order cue) was manipulated between subjects. Third, each participant completed 12-16 runs of the sMGS task, with 16 trials per run, over a 2-2.5 hour session. Hence, the number of trials was increased to 192-256 per subject, aiming for improved modeling performance. Sixteen out of 27 participants completed the quadrant cue and 11 completed the order cue task.

### Oculomotor analysis

The oculomotor preprocessing procedure and analysis were the same as Experiment 1.

All participants completed more than 16 trials per serial position per cue. Out of 256 possible trials in total, on average 11.4% ± 9.1% of trials were excluded across participants for an artifact during the first 500 ms of the response window; 1.9% ± 2.3 % of trials with no defined baseline during the fixation period; 6.5% ± 10.6% of trials with no saccade detected during the response window; 0.2% ± 0.4% of trials with saccade latency shorter than 50 ms; 3.0% ± 3.7% for saccade starting position larger than 2 dva from the center cross; 1.1% ± 1.2% of trials with no fixation detected after the saccade. Participants did not complete 20.2% ± 20.2% of trials on average. After excluding these trials above, 55.8% ± 21.1% of trials were included in the final analysis for the 27 participants, breaking down by 44.7% ± 20.4% of trials as responses to target, 3.7% ± 4.1% of trials as responses to non-target, and 7.4% ± 6.3% of trials as responses outside of all regions of interest.

On average, 36.5 ± 13.8 trials were obtained per serial position (ranging from 20.8 to 60.5 trials) for the quadrant cue task and 34.6 ± 13.6 trials (ranging from 17.3 to 60.3 trials) for the order cue condition. The number of trials included does not significantly differ between cue conditions (*t*(10) = 0.93, p = 0.37).

### Model fitting and statistical analysis

Model fitting and statistical analyses were similar to Experiment 1, except that we conducted a mixed-design ANOVA with serial position as a within-subject factor and with cue condition as a between-subject factor. As the modeled outputs deviated from the normal distribution, we implemented additional nonparametric factorial ANOVAs using the Aligned Rank Transform (ART-ANOVA) (Wobbrock et al., 2011) in R (ARTool) (Kay et al., 2021) on modeled outputs including target rate, swap rate, and guess rate.

### Results Memory-guided saccade error across serial positions

To test for the effects of serial position, cue type, and their interaction on memory-guided saccade endpoint errors, we performed a two-way mixed-design ANOVA (Fig. 4A, B). The analysis showed that errors in the primary saccade significantly varied by serial position (*F*(3, 75) = 4.50, *p* = 0.006, 𝜂_p_^2^ = 0.15) but not by cue type (*F*(1, 25) = 0.42, *p* = 0.52, 𝜂_p_^2^ = 0.02).

**Figure 4.**
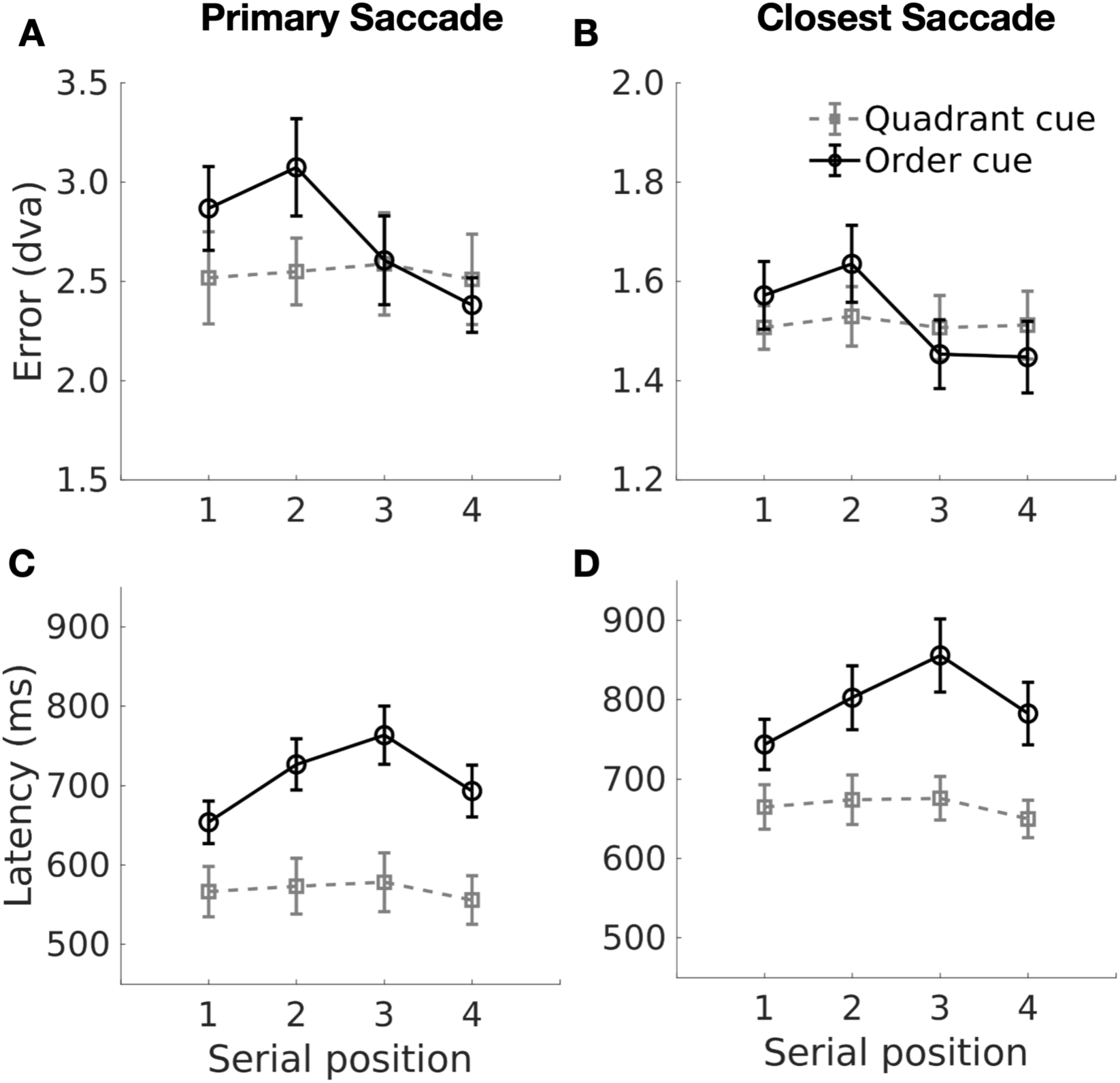
Saccade Endpoint Error and Latency Across Cue Types and Serial Positions in Experiment 2. Figure notations are the same as Fig. 2.

There was also a significant interaction between serial position and cue type (*F*(3, 75) = 4.21, *p* = 0.008, 𝜂_p_^2^ = 0.14). Post hoc analysis revealed that the main effect of serial position was significant for the order cue condition (*F*(3, 30) = 5.20, *p* = 0.005, 𝜂_p_^2^ = 0.34), but not for the quadrant cue condition (*F*(3, 45) = 0.20, *p* = 0.90, 𝜂_p_^2^ = 0.01). In the order cue condition, saccade error decreased linearly with increasing serial position (*t*(30) = -3.25, p = 0.003), with a significant difference between positions two and four (*t*(75) = 0.69, *p* = 0.005, Cohen’s *d* = 1.00).

For the closest saccade to the response target, none of the mixed-design ANOVA effects reached statistical significance, although the effect of serial position approached the significance threshold (*F*(3, 75) = 2.69, *p* = 0.052, 𝜂_p_^2^ = 0.10). A one-way repeated measures ANOVA for each cue condition revealed significant serial-position effects for the order cue (*F*(3, 30) = 3.26, *p* = 0.03, 𝜂_p_^2^ = 0.24) but not for the quadrant cue condition (*F*(3, 45) = 0.09, *p* = 0.97, 𝜂_p_^2^ = 0.006). Specifically, in the order cue condition, errors of the closest saccade to the target decreased linearly with increasing serial positions (*t*(30) = -2.43, p = 0.02), although no post-hoc pairwise comparisons between serial positions were statistically significant.

These findings align with those in Experiment 1, particularly in showing a significant serial-position effect on memory-guided saccade errors during the order cue condition, but not during the quadrant cue condition. However, unlike Experiment 1, no significant differences in saccade endpoint errors were observed between cue types.

### Memory-guided saccade latency across serial positions

Similarly, we conducted mixed-design ANOVAs to examine the effects of serial position, cue type, and their interaction on memory-guided saccade latency (Fig. 4C, D). The latency of both the primary and the closest saccades to the target showed significant main effects of cue type (First saccade: *F*(1, 25) = 8.68, *p* = 0.007, 𝜂_p_^2^ = 0.26; Closest saccade: *F*(1, 25) = 8.59, *p* = 0.007, 𝜂_p_^2^ = 0.26), serial position (First: *F*(3, 75) = 24.65, *p* < 0.001, 𝜂_p_^2^ = 0.50; Closest: *F*(3, 75) = 9.66, *p* < 0.001, 𝜂_p_^2^ = 0.28), and their interaction (First: *F*(3, 75) = 14.21, *p* < 0.001, 𝜂_p_^2^ = 0.36; Closest: *F*(3, 75) = 5.62, *p* = 0.002, 𝜂_p_^2^ = 0.18). Specifically, under the order cue condition, saccade latency varied significantly across serial positions (First: *F*(3, 30) = 30.90, *p* < 0.001, 𝜂_p_^2^ = 0.76; Closest: *F*(3, 30) = 11.16, *p* < 0.001, 𝜂_p_^2^ = 0.53), both linearly (First: *t*(30) = 4.11, *p* < 0.001; Closest: *t*(30) = 2.72, *p* = 0.01) and quadratically (First: *t*(30) = -8.50, *p* < 0.001; Closest: *t*(30) = -4.73, *p* < 0.001). No significant variations in saccade latency across serial positions were observed under the quadrant cue condition for either type of saccade (*p*’s > 0.1).

These results were consistent with Experiment 1, demonstrating a significant effect of cue type on memory-guided saccade latency. Serial-position effects, including both primacy and recency, were significant only during the order cue condition but not the quadrant cue condition.

### Memory precision, swap rate, and transposition errors across serial positions

To examine the sources of serial-position effects, including memory precision, swapping, and guessing, we performed statistical modeling on errors of first memory-guided saccades in Experiment 2 similar to Experiment 1. The standard mixture model best fitted the quadrant-cue data, with the lowest AIC (894.4 ± 288.1) and BIC (866.8 ± 298.7) values compared to other models. For the order cue data, the swap model had the lowest BIC (862.6 ± 295.5), while the swap plus systematic bias model had the lowest AIC (852.4 ± 293.8), indicating more item swapping in the order cue condition (Table 2).

**Table 2.** ANOVA on Modeled Outputs for Behavioral Experiment 2. ANOVA: Mixed design ANOVA; ART-ANOVA: Aligned Rank Transform ANOVA (Wobbrock et al., 2011)

|  |  | Cue | Serial position | Cue x Serial position |
| --- | --- | --- | --- | --- |
| Memory imprecision | ANOVA | $F(1,25) = 0.57, p = 0.46, \eta_p^2 = 0.02$ | $F(3,75) = 1.02, p = 0.39, \eta_p^2 = 0.04$ | $F(3,75) = 0.28, p = 0.84, \eta_p^2 = 0.01$ |
| | ART-ANOVA | $F(1,25) = 0.48, p = 0.50$ | $F(3,75) = 1.92, p = 0.13$ | $F(3,75) = 0.16, p = 0.92$ |
| Target rate | ANOVA | $F(1,25) = 3.02, p = 0.10, \eta_p^2 = 0.11$ | $F(3,75) = 1.76, p = 0.16, \eta_p^2 = 0.07$ | $F(3,75) = 2.61, p = 0.06, \eta_p^2 = 0.10$ |
|  | ART-ANOVA | **$F(1,25) = 18.44, p < 0.001$** | **$F(3,75) = 2.82, p = 0.04$** | **$F(3,75) = 4.61, p = 0.005$** |
| Swapping rate | ANOVA | $F(1,25) = 3.81, p = 0.06, \eta_p^2 = 0.13$ | $F(2.1, 53.5) = 1.94, p = 0.13, \eta_p^2 = 0.07$ | $F(2.1, 53.5) = 2.20, p = 0.12, \eta_p^2 = 0.08$ |
|  | ART-ANOVA | **$F(1,25) = 28.44, p < 0.001$** | **$F(3,75) = 6.89, p < 0.001$** | **$F(3,75) = 8.41, p < 0.001$** |
| Guessing rate | ANOVA | $F(1,25) = 0.98, p = 0.33, \eta_p^2 = 0.04$ | $F(1.7, 43.1) = 0.62, p = 0.61, \eta_p^2 = 0.02$ | $F(1.7, 43.1) = 0.33, p = 0.81, \eta_p^2 = 0.01$ |
| | ART-ANOVA | $F(1,25) = 0.49, p = 0.49$ | $F(3,75) = 0.48, p = 0.69$ | $F(3,75) = 0.76, p = 0.51$ |

To compare memory precision, swapping, and guessing rates across cue conditions, we use the swap model to fit the first saccade endpoint distributions for both cue conditions. Following the model estimation, we conducted mixed-design ANOVAs with serial position as a within-participant factor and cue type as a between-participant factor on the parameter estimates. None of the ANOVA effects were statistically significant for any modeled outputs, including memory imprecision, target rate, swapping rate, and guessing rate (Table 3). However, the model outputs, particularly target rate, swapping rate, and guessing rate, significantly deviated from a normal distribution (Shapiro-Wilk test, *p*’s < 0.001 for quadrant cue). Therefore, we conducted an additional mixed-design ART-ANOVA on these parameters.

The mixed-design ART-ANOVA revealed significant effects of cue type, serial position, and their interaction on swapping rate (Fig. 5B) and target rate (Fig. 5C), but not on memory imprecision (Fig. 5A) and guessing rate (Fig. 5D; Table 3). Specifically, for swapping rate, there were significant effects of cue type (*F*(1,25) = 28.44, *p* < 0.001), serial position (*F*(3,75) = 6.89, *p* < 0.001), and their interaction (*F*(3,75) = 8.41, *p* < 0.001).

**Figure 5.**
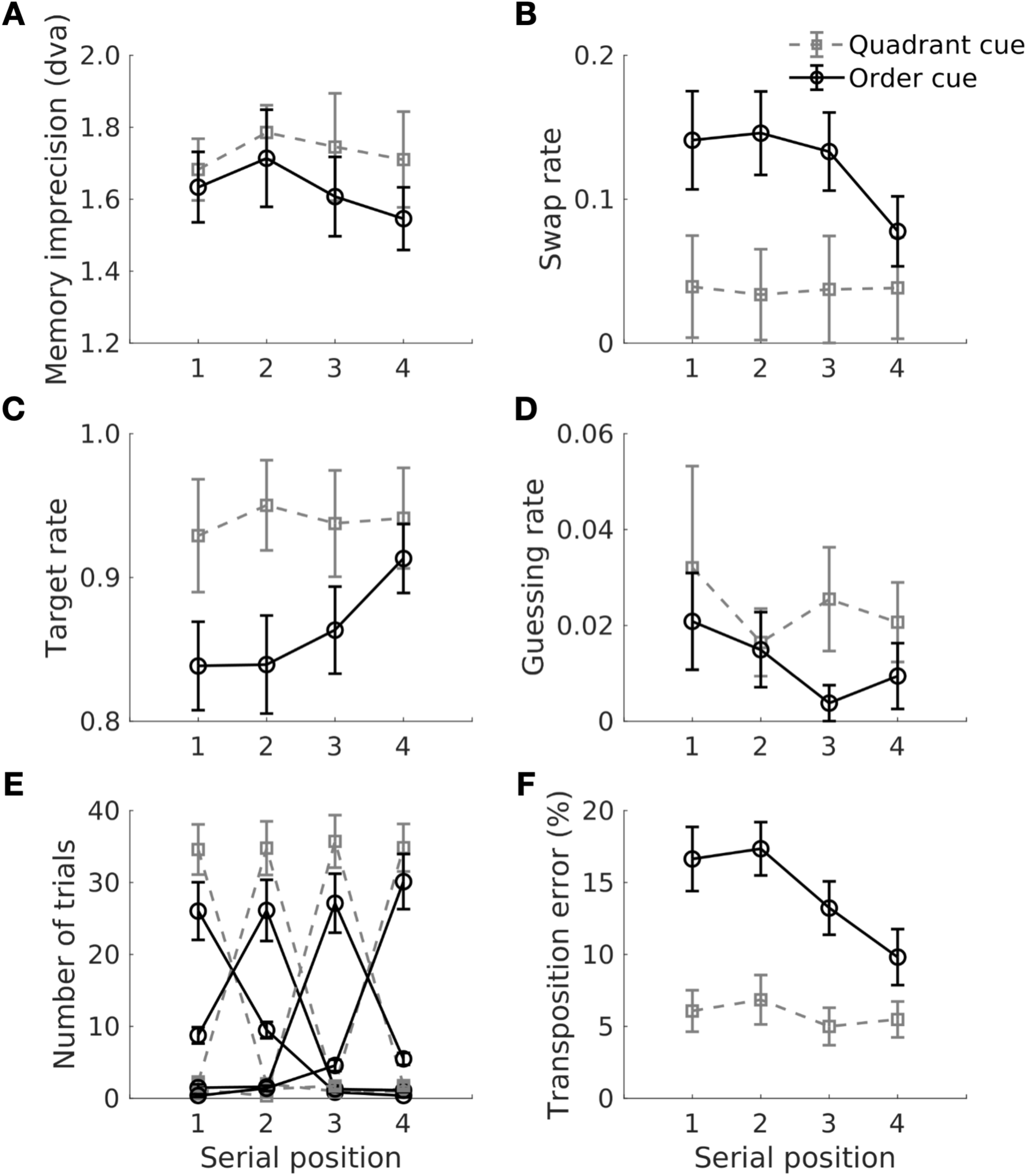
The Swap Model Results and Transposition Error of Memory-Guided Saccades in Experiment 2. Figure notations are the same as Fig. 3.

Similar to Experiment 1, the transposition error of the first saccade (Table 3; Fig. 5E, F) varied significantly by cue type (*F*(1,25) = 22.76, *p* < 0.001, 𝜂_p_^2^ = 0.48) and serial position (*F*(3,75) = 4.87, *p* = 0.004, 𝜂_p_^2^ = 0.16). The interaction between serial position and cue type approached significance (*F*(3, 75) = 2.49, *p* = 0.07, 𝜂_p_^2^ = 0.09). Specifically, transposition error was significantly higher during the order cue than the quadrant cue condition (*t*(25) = 4.77, *p* < 0.001, Cohen’s *d* = 1.38). Significant serial-position effects on transposition error were observed for the order cue (*F*(3, 30) = 3.20, *p* = 0.04, 𝜂_p_^2^ =0.24) with a linear trend (*t*(30) = -2.84, *p* = 0.008) but not for the quadrant cue condition (*F*(3, 45) = 1.04, *p* = 0.38, 𝜂_p_^2^ =0.07).

Collectively, Experiment 2 yielded results largely consistent with Experiment 1, despite differences in design and sample. Significant serial-position effects on endpoint error, latency, transposition error, and modeled swap rate of memory-guided saccades were observed in the order cue condition but not the quadrant cue condition, although the primacy effects were weaker compared to Experiment 1. Similar to Experiment 1, serial-position effects on modeled memory imprecision were not statistically significant. Unlike Experiment 1, which showed a significant cue effect on all saccade measures, Experiment 2 only revealed higher transposition error and modeled swap rate during the order cue condition than the quadrant cue condition. This weaker cue-related effect in Experiment 2 may be attributed to the between-participant manipulation of cue type, in contrast to the within-participant manipulation in Experiment 1.

## Discussion

In two experiments, we showed that explicit serial-position demand (order cue) led to greater recall error and latency in memory-guided saccades and significant serial-position effects, compared to when serial position was implicit and not required for task completion (quadrant cue) (Figs. 2, 4). Specifically, saccades to the intermediate items in a sequence were less accurate and took longer to initiate than those to more recent items (recency effects, observed in both experiments) and earlier items (primacy effects, more prominent in Experiment 1). Statistical modeling revealed that the order-cue data was best fit by a swap model, whereas the quadrant-cue data was best fit by a non-swap model with no clear serial-position modulation of the swap rate (Figs. 3, 5). Overall, these results suggest that misbinding between spatial location and serial position is a key factor leading to reduced memory-guided saccade precision for intermediate items in a working memory sequence, resulting in the observed primacy and recency effects, particularly when order is required for task completion.

### Feature Misbinding as a Major Source of Serial-Position Effects

Although serial-position effects have been observed for decades in short-term memory (Hurlstone et al., 2014), their underlying sources remain debated. Previous studies have proposed several explanations, including inter-item interference during memory retrieval (Brown et al., 2000; McAteer et al., 2023) and uneven allocation of memory resources across items during retention (Gorgoraptis et al., 2011; McAteer et al., 2023; Udale et al., 2022; Zokaei et al., 2011). Intermediate items may suffer from stronger proactive and retroactive interference, reducing memory precision and increasing the likelihood of feature misbinding.

Our data highlight feature misbinding as a key driver of serial-position effects in visuospatial working memory. Both experiments revealed robust serial-position effects on memory-guided saccades under explicit, but not implicit, order demand. Statistical modeling confirmed that these effects were associated with increased target-non-target swaps, rather than decreased precision in target representation across items in different serial positions, indicating that misbinding between spatial and ordinal features underlies the observed primacy and recency effects.

Our results align with earlier studies showing serial-position modulation on item swapping during delayed recall of motion (Zokaei et al., 2011; Exp. 1) and orientation (Gorgoraptis et al., 2011; Exp. 1) at a similar set size. Specifically, we observed both primacy and recency effects on swap rates, consistent with previous findings. However, our findings diverge from McAteer et al. (2023), which did not observe significant serial-position effects on item swapping in a spatial working memory task. This discrepancy may stem from differences in sample size, as McAteer et al.’s study involved only 10 participants, which may have been underpowered.

Extending previous research focused on visual or visual-spatial feature misbinding, we demonstrated that ordinal-spatial misbinding significantly contributes to serial-position effects. This finding echoes neural evidence showing that lateral prefrontal neurons exhibit selectivity for combinations of spatial and visual features (Rainer et al., 1998; Rao et al., 1997), visual and ordinal features (Ninokura et al., 2004), and spatial and ordinal features (Carpenter et al., 2018). These features may be bound via similar mechanisms such as multiplicative gain modulation (Botvinick & Watanabe, 2007), neural tensor products (Smolensky, 1990; Xie et al., 2022), or temporal synchronization (Parto Dezfouli et al., 2021; Siegel et al., 2009). The hippocampus may also support feature binding via cognitive mapping (Nieh et al., 2021). Both the prefrontal cortex and hippocampus may support the neural substrates for binding diverse features required for goal-directed behaviors. Further research is needed to elucidate the neural mechanisms underlying the serial-position effects on the feature-binding process.

### Serial-Position Effects on Memory Precision

Contrary to our hypothesis, we did not observe robust serial-position effects on spatial memory precision beyond location-serial-position misbinding. In the quadrant cue condition, where order demand was implicit or not required for task completion, there were no significant effects of serial position on memory-guided saccade error or latency. Moreover, closest saccades to the target, reflecting refined memory responses, varied even less across serial positions compared to first saccades and did not consistently reach statistical significance across experiments. Similarly, modeled memory precision did not significantly differ by serial position, even under explicit order demand.

These findings align with studies showing no significant serial-position effects on spatial or motion working memory precision (McAteer et al., 2023, Exp. 3; Zokaei et al., 2011, Exp. 1), but contradict with studies that found significant effects on orientation memory precision (Gorgoraptis et al., 2011, Exp. 1). It is possible that serial-position effects on memory precision are more pronounced for visual features than for spatial features.

Moreover, prior studies investigating working memory precision across serial positions often incorporated additional visual features (e.g., color) as contextual cues, thereby introducing additional feature-binding load and confounding measurements of memory precision. In contrast, our quadrant cue design required no explicit feature binding, potentially providing a more accurate assessment of the effects of serial position on spatial memory precision. However, it is also possible that the spatial nature of the quadrant cue minimized serial-position effects, producing more accurate but less differentiated responses across serial positions.

Another potential explanation involves participants’ knowledge of task structure. In our design, the number of items was fixed and all items were presented and probed with equal probability. This task structure may have encouraged participants to adopt a strategy of evenly distributing memory resources across all items, leading to more uniform memory precision across serial positions compared to tasks with variable set sizes (McAteer et al., 2023).

Nevertheless, these results suggest that serial-position effects under a fixed set size may not involve trading representation precision across items but are instead linked to more item swapping for intermediate items. For example, Gorgoraptis et al. (2011) found that while both memory precision and misbinding rates varied by serial position, only misbinding rates differed significantly between sequential and simultaneous presentation conditions (Exp. 1 vs. 2), highlighting the central role of feature misbinding in serial-order processing. Conversely, changes in memory precision are more commonly associated with variations in set size rather than serial position. Memory precision has been consistently shown to decline with increasing set size, either for all items (Gorgoraptis et al., 2011) or specifically for the most recent item in a sequence (Zokaei et al., 2011).

### Saccade Metrics and Cognitive Components of Spatial Working Memory

Our eye-tracking approach offered new insights into the relationship between saccade metrics and cognitive components of spatial working memory. We observed a strong interaction between serial position and cue condition on first saccades made during the response window, such that the error of the first saccade varied significantly by serial position during the order cue but not quadrant cue. In contrast, the closest saccades to the response targets showed similar serial-position modulation across cue conditions, particularly when the cue was manipulated within participants (Fig. 2). This distinction mirrors the differences between modeled swap rate and memory precision (Fig. 3). These findings suggest that different types of saccades may reflect different cognitive components of spatial working memory. Whereas the primary memory-guided saccades may represent a crude memory representation involving item swapping, closest saccades may reflect a more refined memory representation of the target.

Besides different types of saccade, our study also revealed novel findings about different measures of saccades including error and latency. While previous studies have primarily focused on examining response accuracy across serial positions, we found a strong primacy and recency effect on saccade latency during the order cue condition but this was not evident during the quadrant cue condition. These results suggest that saccade latency might be a robust indicator of serial-order processing. Specifically, the serial-position curve of latency closely resembles that of endpoint error, transposition error, and modeled swap rate of first saccades, indicating that latency is a sensitive measure that closely reflects the misbinding between spatial location and serial position.

### Conclusion and limitations

One limitation with our design is the limited spatiotemporal patterns, always presenting stimuli from left to right or vice versa. Such order regularities may result in specific behavioral strategies, such chunking, which could influence task performance (Rosenbaum et al., 1983). Previous studies suggest that different relational structures of sequences affect response accuracy in delayed sequence reproduction tasks (Zhang et al., 2022). Future studies could explore whether different order patterns, such as regularity or predictability, impact serial-position effects.

In conclusion, our study demonstrates that order demand dynamically distributes working memory resources across serial positions. When serial position is task-irrelevant, items are represented with similar accuracy. However, when order is explicitly required, intermediate items are less precisely represented than earlier and later items. Notably, these serial-position effects are primarily driven by misbinding between spatial location and serial position, rather than variation in spatial memory precision.

