## Supplemental Figures for "Feature Misbinding Underlying Serial-Order Effects of Visuospatial Working Memory"

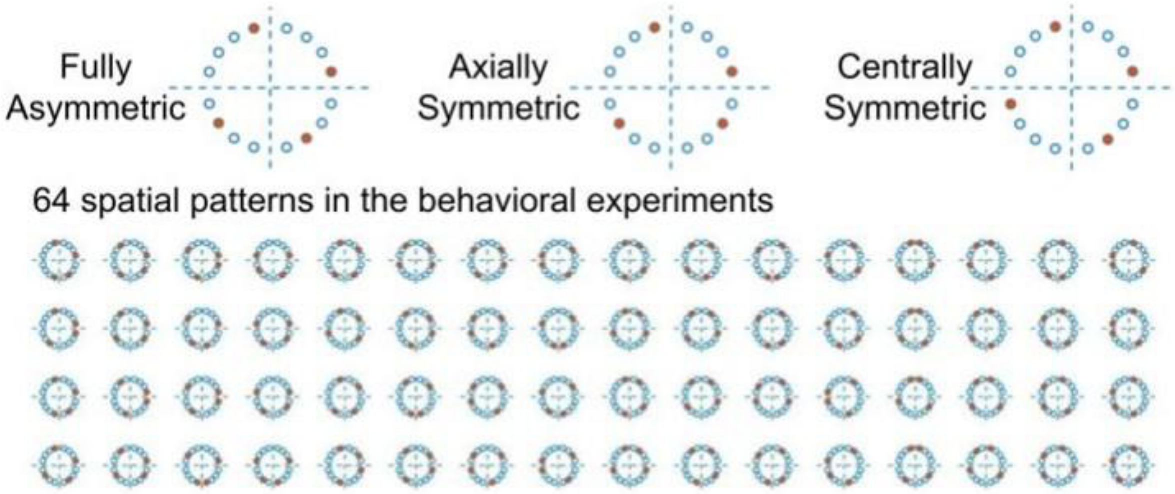


**Figure S1. Study 1 Spatial Patterns of sMGS Task Stimuli**. To focus on studying the effects of serial position rather than differences between sequence patterns, we constructed an array of spatial sequences that covered all four quadrants and crossed through the two sides equally. We initially generated 256 possible sets of locations, each with 16 distinct spatial patterns. In each set, all 16 polar angles appeared an equal number of times. We then selected patterns that were fully asymmetric or had only one pair of axially symmetric angles or one pair of centrally symmetric angles, resulting in 168 viable sets out of the original 256. We further randomly selected four sets out of these 168 sets 1000 times and chose one selection that exhibited minimal overlapping of spatial patterns or maximum pattern dissimilarity based on Procrustes analysis. The final set of sequences comprised 4 sets of 16 spatial patterns each as shown in the figure, and they were used in the 4 experimental runs (16 trials each). Among these 64 patterns used, there are 54 unique spatial patterns, with ten of these patterns repeated once.

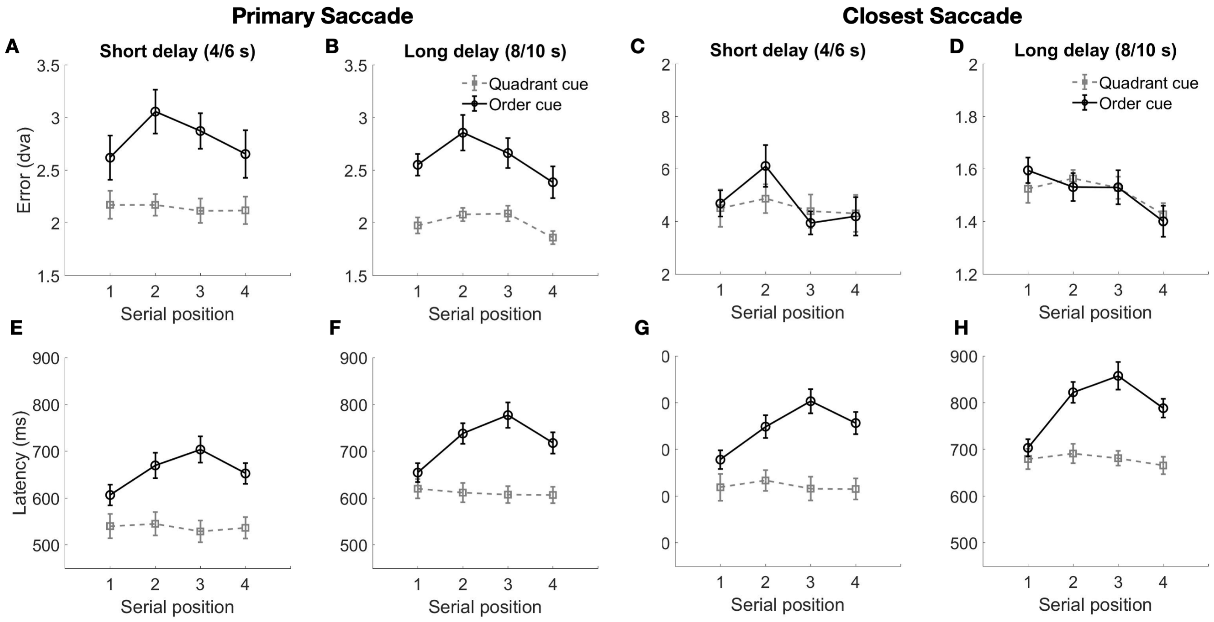


**Figure S2. Study 1 Effects of Delay Duration on Saccade Error and Latency in Experiment 1**. Delay was manipulated between participants: 30 participants completed the task with a short delay (4/6 s) and 35 participants completed the task with a long delay (8/10 s). Top row (A-D) shows the endpoint error plots. Bottom row (E-G) shows the latency plots. Columns from left to right display the plots of primary saccades for the short (A, E) and long (B, F) delay conditions, and closest saccade to target for the short (C, G) and long (D, H) delay. Y axis represents the saccade measures and X axis represents the serial position of the target as first, second, third, or fourth item. Different cue types are plotted as different lines: Quadrant cue is plotted in gray and rectangle marker; Order cue is plotted in black and round marker. Cue conditions were manipulated within participants. Three-way mixed design ANOVAs with serial position and cue as within-participant factor and delay as between-participant factor did not reveal significant effects of delay, except for the latency of the first saccades ($F(1,63) = 5.59$, $p = 0.02$, $\eta_p^2 = 0.06$). Particularly, first saccade latency was significantly longer for long delay than short delay ($t(63) =$

2.36, $p = 0.02$, Cohen's $d = 0.53$). None of the interaction effects with delay reached statistical significance.
